# Glycolytic metabolism of pathogenic T cells enables early detection of GvHD by ^13^C-MRI

**DOI:** 10.1101/2020.03.16.984609

**Authors:** Julian C. Assmann, Don E. Farthing, Keita Saito, Natella Maglakelidze, Brittany Oliver, Kathrynne A. Warrick, Carole Sourbier, Christopher J. Ricketts, Thomas J. Meyer, Steven Z. Pavletic, W. Marston Linehan, Murali C. Krishna, Ronald E. Gress, Nataliya P. Buxbaum

## Abstract

Graft-versus-host disease (GvHD) is a prominent barrier to allogeneic hematopoietic stem cell transplantation (HSCT). Definitive diagnosis of GvHD is invasive and biopsies of involved tissues pose a high risk of bleeding and infection. Our previous studies in a chronic GvHD mouse model demonstrated that alloreactive CD4^+^ T cells are distributed to target organs ahead of overt symptoms, meanwhile CD4^+^ T cell activation is tied to increased glycolysis. Thus, we hypothesized that metabolic imaging of glycolysis would allow non-invasive detection of insipient GvHD in target organs infiltrated by glycolytic effector memory CD4^+^ T cells. We metabolically characterized CD4^+^ T cell subsets on day 14 post-transplant before the onset of chronic GvHD in a pre-clinical mouse model and performed ^13^C hyperpolarized magnetic resonance imaging (MRI) to quantify glycolytic activity in the liver of mice over the course of the disease. Intracellular metabolic screening and *ex vivo* metabolic profiling of CD4^+^ T cell subsets at day 14 confirmed that activated CD4^+^ T cells were highly glycolytic. Concurrently, hyperpolarized ^13^C-pyruvate MRI of the liver showed high conversion of pyruvate to lactate, indicative of increased glycolytic activity, that distinguished allogeneic from syngeneic HSCT recipients prior to the development of overt chronic GvHD. Furthermore, single cell sequencing of T cells in patients undergoing allogeneic HSCT indicated that similar metabolic changes may play a role in acute GvHD, providing a rationale for testing this imaging approach in the clinical post-HSCT setting. Our imaging approach is amenable to clinical translation and may allow early, non-invasive diagnosis of GvHD.

## Introduction

Allogeneic hematopoietic stem cell transplantation (AHSCT) is a curative therapy for patients with aggressive hematologic malignancies, non-malignant hematologic diseases, and a consolidation therapy for patients receiving chimeric antigen receptor T cells. Acute and chronic graft-versus-host disease (GvHD) following AHSCT are prevalent, morbid, and may share pathophysiologic features. Acute GvHD (aGvHD) is the most significant risk factor for the development of chronic GvHD (cGvHD) and jointly they represent a major barrier to successful AHSCT^1–3^. Additionally, GvHD of the liver and the gastrointestinal tract pose a particular diagnostic challenge^4,5^. Biopsies of these tissues are invasive, associated with serious bleeding and infection risks for the AHSCT recipient, and do not guide treatment selection. Responses to therapy are also difficult to assess and are being ascertained by tabulating and tracking the severity of clinical signs and symptoms^2,4–7^. Non-invasive objective approaches to diagnose and monitor GvHD are needed^8^.

T cells are the primary mediators of GvHD^2^. Therefore, therapies and prevention strategies for GvHD have targeted T cells^9,10^, their subsets^11,12^, differentiation^13,14^, survival^15^, and milieu, including antigen presenting cells^16,17^ and cytokines^18,19^. T cell activation is an early event in GvHD pathogenesis and thus an attractive target for early diagnosis and intervention. Antigen-driven CD4^+^ T cell activation has been directly tied to increased rates of glycolysis and increased glycolytic flux is essential for CD4^+^ T cell effector function^20–22^. We previously showed that target organs of cGvHD are infiltrated by pathogenic alloreactive CD4^+^ T effector memory (T_em_) cells in advance of overt disease^23^. Thus, we hypothesized that metabolic tracing of a glycolytic metabolite in GvHD target organs may allow non-invasive detection of GvHD *in vivo.*

Metabolic imaging provides important biological insights when combined with anatomical imaging. Currently, the most widely used metabolic imaging technique, fluorodeoxyglucose (FDG)-positron emission tomography (PET), detects increased glucose uptake but not its downstream metabolites. As a result, FDG-PET is unable to distinguish increases in glycolytic flux from increases in oxidative phosphorylation^24^. To overcome this limitation ^13^C-labeled metabolites detected by spectroscopic magnetic resonance imaging (MRI) have become increasingly used for functional imaging primarily in the setting of malignancy^25–30^, and more recently in other inflammatory conditions^31,32^. *Ex vivo* dynamic nuclear hyperpolarization (DNP) of the ^13^C-pyruvate tracer prior to infusion has improved the sensitivity of its MRI detection to allow real-time spectroscopic measurement of *in vivo* glycolytic activity within tissues in pre-clinical and clinical studies^33,34^. An additional benefit of this methodology is the lack of exposure to radioactivity from the tracer and/or scanner.

We hypothesized that early post-transplant antigen stimulated T cells would have elevated glycolytic metabolism, which could allow *in vivo* detection of GvHD via metabolic imaging of target organs infiltrated by these cells. Thus, we applied hyperpolarized ^13^C-pyruvate MRI to an established minor histocompatibility antigen-mismatched GvHD model^23,35–38^ to evaluate whether increased glycolysis within a target organ could be detected ahead of overt clinical disease manifestations. Furthermore, we ascertained whether similar metabolic patterns would be present in circulating CD4^+^ T cells in patients prior to onset of aGvHD clinical symptoms, which would further support the potential for metabolic imaging to non-invasively detect insipient GvHD in patients undergoing AHSCT.

## Methods

### Mice

Female BALB/cAnNCr (H-2^d^, #555) mice were purchased from Charles River (Wilmington, MA, USA). Age-matched female B10.D2 (H-2^d^, #000463) mice were acquired from Jackson Laboratory (Bar Harbor, ME, USA). All mice were acclimatized for at least 4 weeks before transplantations and kept on a 12h/12h light-dark cycle with unrestricted access to food and water in a specific pathogen free environment. All animal studies were reviewed and approved by the NCI Animal Care and Use Committee.

### GvHD

Recipient female mice at 12-13 weeks of age were conditioned with 850 cGy total body irradiation (TBI) delivered in two doses 3 hours apart on day −1. Age-matched BALB/c mice served as donor for matched, syngeneic transplants whereas B10.D2 mice were used for allogeneic transplants. Recipients were reconstituted with 15 million unfractionated splenocytes and 8 million bone marrow cells injected via tail vein on day 0. The injection buffer contained gentamicin (100 μg/ml), however no additional antibiotics were administered posttransplant. Animals were weighed and cGvHD scoring was performed twice weekly using a previously described clinical scoring system^39^.

### Organ collection and preparation

#### Spleen

Single-cell suspensions were obtained by mechanically disrupting harvested spleens and filtering through 70-μm mesh filters (Thermo Fisher Scientific). Ammoniumchloride potassium (ACK) lysis was performed to remove erythrocytes prior to cell counting.

#### Bone marrow

*Cells* were flushed from femur and tibia of donor mice and filtered through 70-μm mesh filters. Erythrocytes were lysed using ACK.

#### Blood

Circulating immune cells were isolated from blood drawn via cardiac puncture and subsequently filtered and lysed as described above.

### Flow cytometry and fluorescence-activated cell sorting (FACS)

One million to two million live cells were aliquoted for flow cytometry staining. For fluorescence-activated cell sorting (FACS), samples from multiple mice were pooled for each cohort (syngeneic recipients and allogeneic recipients). A list of surface antibodies is provided in the Table 1. mCD1d (PBS-57) tetramer was obtained from the NIH Tetramer Core Facility to allow exclusion of NKT cells from CD4^+^ T cells for FACS and flow cytometry phenotyping. Pooled samples for each cohort underwent FACS using BD Influx achieving 95% purity. Individual mouse sample flow cytometry measurements were obtained on a BD LSR II or Fortessa (Becton Dickinson, USA), and analyzed using FlowJo 9.7.6 Software (BD). FACS-purified samples were collected into PBS buffer containing 2% (v/v) BSA. Sorted T cell subsets were further processed for use in metabolomics analysis, extracellular flux analysis and RNA-sequencing as described below.

**Table 1.** Flow cytrometry antibodies

| **<u>Antigen</u>** | **<u>Clone</u>** | **<u>Manufacturer</u>** |
| --- | --- | --- |
| CD4 | GK1.5 | eBioscience |
| CD8a | 5H10 | Invitrogen |
| CD69 | H1.2F3 | Biolegend |
| CD44 | IM7 | Biolegend |
| CCR7 (CD197) | 4B12 | Biolegend |
| CD1d | PBS57 tetramer | NIH Tetramer Core facility |
| CD16/32 (Fc block) | 2.4G2 | BD Biosciences |
| Live/Dead Aqua viability dye |  | Invitrogen |

### Metabolomics Analysis

FACS-purified T cell subsets (~500,000 cells) were collected directly into a 15 mL polypropylene centrifuge tube and processed immediately. The cells were centrifuged (~314 × g, 5 min, room temperature (RT)), the supernatant was discarded, and the pellet was resuspended in 10 mL aqueous mannitol (5%, w/v) by gentle mixing. The cell suspension was centrifuged (~314 × g, 5 min, RT) and the supernatant (aqueous mannitol) was discarded. The pellet was then resuspended in deionized water (50 μL) and centrifugation (~314 × g, 5 min, RT) was repeated. Approximately 100-150 μL of the supernatant was discarded followed by addition of 133.3 μL methanol (mixed for 30 sec) to quench intracellular enzymes. The working internal standard solution (10 μM) used for mass spectrometry (MS) analysis was added (91.7 μL), mixed for 30 sec and samples were stored at −80 °C. The processed T cell samples were pooled and quantified relatively utilizing capillary electrophoresis time-of-flight mass spectrometry (CE-TOF-MS, Agilent Technologies, Waldbronn, Germany) with analysis performed by Human Metabolome Technologies, Inc. (HMT, Japan). The samples were thawed, combined, vortexed and 1,100 μL was transferred to a 1.5 mL microcentrifuge tube and centrifuged (2,300 × g, 5 min, 4 °C). Two ~500 μL aliquots were transferred to 5 kDa MWCO centrifugal filter units (0.5 mL, Millipore Sigma, Burlington, MA, USA) and centrifuged (9,100 × g, 4 °C) until sample filtration was complete (~2-5 hrs.). The sample filtrates were transferred to 1.5 mL polypropylene microcentrifuge tubes and vacuum dried to completeness at room temperature using a Savant DNA 110 Speed Vac^®^ (Savant Instruments, Inc., Farmingdale, NY, USA). The sample residue containing ~ 3 million T cells for MS analysis was capped and stored at −80 °C. After shipment to HMT, the sample residues were reconstituted with 50 μL deionized water, mixed and ~100 nL injected for CE-TOF-MS analysis.

### Metabolic flux assay

Mitochondrial function and glycolysis were evaluated using a Seahorse XFe96 Bioanalyzer (Agilent) per the manufacturer instructions. Subsets of CD4^+^ T cells were FACS-purified and plated into 96-well plates coated with CellTak (Corning, #354240) with a seeding density of 100,000 live cells per well and a minimum of 8 wells per cell subset. Oligomycin (1 μM), 2-[[4-(trifluoromethoxy)phenyl]hydrazinylidene]propanedinitrile (FCCP, 0.5 μM) and Rotenone/antimycin (1 μM/ 1 μM) were loaded into the injection ports, injected as indicated in the figures and used to measure the oxygen consumption rate and extracellular acidification rate with a total plate run time of up to 140 minutes.

### RNA extraction and quantification

RNA was extracted from sorted cell populations using the RNeasy^®^ Plus Mini Kit following the manufacturers protocol for purification of total RNA from animal cells (Qiagen). In brief, the cells were lysed in Buffer RLT Plus with β-ME, homogenized with a QIAshredder spin column, and the resultant lysate was run through a gDNA Eliminator spin column to remove genomic DNA. RNA was extracted by passing the remaining lysate through a RNeasy spin column and, following washes, total RNA was resuspended in 30 μl of RNase-free water. Measurement of RNA concentration and quality control was performed using a 2100 Bioanalyzer instrument with the standard manufacturers protocol (Agilent).

### Bulk RNA Library and Sequencing Method

RNA-Seq libraries were constructed from 0.1-0.6 μg total RNA. The HyperPrep RNA-Seq Kit with Riboerase (Kapa Biosystems/Roche) was used according to manufacturer’s instructions to deplete rRNA and then construct libraries for sequencing on Illumina platforms. Amplification was performed using 10 or 12 cycles which was optimized for the input amounts and to minimize the chance of over-amplification. Unique barcode adapters were applied to each library. Libraries were pooled in equimolar ratio for sequencing. The pooled libraries were sequenced on multiple lanes of a HiSeq 4000 to achieve a minimum of 65 million 75 base read pairs. The data was processed using RTA version 1.18.54 and CASAVA 1.8.2.

### Bulk RNA sequencing

RNA-seq NGS-datasets were processed using the CCBR Pipeliner utility (https://github.com/CCBR/Pipeliner). Briefly, reads were trimmed of low-quality bases and adapter sequences were removed using Trimmomatic v0.33^40^. Mapping of reads to the Gencode mm10 mouse reference genome (M12 release) was performed using STAR v2.5.2b in 2-pass mode^41^. Then, RSEM v1.3.0 was used to quantify gene-level expression, with counts normalized to library size as counts-per-million^42^. Finally, limma-voom v3.34.5 was used for quantile normalization and differential expression^43^.

Data for the mitochondrial genome-encoded genes for the electron transport chain were not evaluated and the genes for complex II (succinate dehydrogenase) were selected as part of the Krebs cycle genes. For the RNAseq analysis, the CPM (counts per million) with TMM (trimmed mean of M values) normalization data was used to assess differences in gene expression between samples. For each gene, the expression of the T_em_ cells was compared to the expression of phenotypically naïve T cells (T_n_) and expressed as a ratio (T_em_ / T_n_).

### Hyperpolarized ^13^C-Pyruvate MRSI

MRI scans were performed on a 3T scanner controlled with EasyScan and PowerScan (MR Solutions, Acton, MA). Mice were put in a 35 mm ^13^C-^1^H saddle coil (custom built in-house). T2-weighted anatomical images were obtained using a fast spin echo sequence (FSET2) with TE of 13 ms, TR of 2,500 ms, 8 slices, 2 mm thickness and a resolution of 0.25 × 0.25 mm. [1-^13^C] pyruvic acid (30 μL, Cambridge Isotope Laboratories, Tewksbury, MA) containing 15 mM OX063 and 2.5 mM gadolinium chelate ProHance (Bracco Diagnostics, Milano, Italy) was hyperpolarized at 3.35 T and 1.45 K using a Hypersense DNP polarizer (Oxford Instruments, Abingdon, UK) for 40–60 min. The hyperpolarized [1-^13^C]pyruvic acid was rapidly dissolved in 4.5 mL of a superheated alkaline buffer comprising 50 mM Tris, 60 mM NaOH, and 100 mg/L ethylendiaminetetraacetic acid to a final concentration of 96 mmol/L, and was intravenously injected into the mouse through a catheter placed in the tail vein (12 μL/g body weight). ^13^C twodimensional spectroscopic images were acquired 25 s after the beginning of the [1-^13^C]pyruvate solution injection, with a 32 × 32 mm of field of view in an 8 mm coronal slice through the body, a matrix size of 16 × 16, spectral width of 3330 Hz, repetition time of 85 ms, and an excitation pulse with a flip angle of 20°. The total time required to acquire each image was 22 s.

### Lactic acid measurement

Mouse plasma L-lactate levels were quantified via mass spectrometry in mandibular blood samples at indicated timepoints. For quantitatively measuring L-lactic acid levels in mouse plasma, we used a Waters Acquity H-Class UPLC system coupled to a Waters Xevo ESI-MS/MS. As mouse blood volume is limited, we developed a MS method that utilized only 25 to 50 μL of mouse plasma, which was collected using a fluoride/oxalate microtainer tube. Briefly, the mouse plasma and internal standard (D3-lactic acid) were added to a 3K MWCO filter (Amicon^®^, 0.5 mL) and centrifuged (14,000 × g, 30 min, 20 °C). The filtrate was transferred to a Waters autosampler microvial (deactivated) and 2 μL was injected for MS analysis. The 3K MWCO filter provided clean extracts for UPLC HILIC chromatography and the electrospray ionization ESI (-) SIR (selected ion recording) mode of MS analysis. For HILIC plasma component separation, we used a linear gradient consisting of aqueous ammonium acetate (26 mM) and acetonitrile, flow rate of 0.3 mL/min, and column temperature of 30 °C. The HILIC chromatography provided good resolution of the plasma components and excellent peak shape for L-lactic acid and the internal standard. The quantitative SIR MS method utilized five calibration standards (L-lactic acid, 0.28-5.6 mM) and a deuterated internal standard (D3-lactic acid, 1.1 mM). The UPLC-ESI (-) SIR-MS method demonstrated good sensitivity (MS response), specificity and linearity using a quadratic fit calibration model for quantifying L-lactic acid.

### scRNAseq patient characteristics

We screened annotated clinical data from a GvHD clinical study performed at the NIH Clinical Center (NCT00520130). Two patients that matched our criteria were identified. One patient (“pre-GvHD”) received an 8/8 matched unrelated donor (MUD) peripheral blood hematopoietic stem cell transplant (PBHSCT) for CML in remission. The other (“No GvHD”) received an 8/8 MUD PBHSCT for relapsed mantle cell lymphoma in remission. Both patients received reduced intensity conditioning with fludarabine and cyclophosphamide and were receiving methotrexate, tacrolimus and sirolimus for GvHD prophylaxis at the time of PBMC collection. The blood samples for both patients were collected on post-transplant day +30. The patient “pre-GvHD” developed an erythematous rash and nausea on post-transplant day +35, with upper GI biopsy on day +37 consistent with GvHD, i.e. epithelial apoptosis in the duodenum, stomach antrum and body (CMV and *H.pylori* negative), while patient “No GvHD” did not have symptoms of acute GvHD, but did have ongoing CMV retinitis that preceded HSCT and for which systemic anti-virals had been discontinued on day −1 with once weekly intravitreous anti-viral injection ongoing at the time of sample collection. Since acute GvHD significantly increases chances of developing chronic GvHD we decided to exclude patients that may have had sub-clinical undiagnosed acute that later went on to develop cGvHD. Thus, we selected our “No GvHD” patient based on the fact that at 3 years post HSCT this patient showed no signs of cGvHD.

### scRNAseq sample preparation analysis

Frozen PBMC samples were thawed rapidly for 90 s at 37 °C, slowly resuspended in RPMI supplemented with 5% fetal bovine serum and immediately put on ice. Cells were spun at 300 × g for 10 min, resuspended in 1 ml RBC lysis buffer (Thermo Fisher Scientific, #00-4333-57) and incubated at room temperature for 5 min. Following lysis, cells were washed twice to remove ambient contaminating mRNA, reduced to a volume of 100 μl and viability was assessed using a LunaFL cell counter (Logos Biosystems). Viability for both samples was >90%. To profile the 5’ single cell gene expression, single cell suspensions were then loaded onto a Chromium Single Cell Chip (10X Genomics) with a recovery target of 6,000 cells per lane according to the manufacturer’s instructions. All subsequent steps of cDNA generation, library preparation and quality control were performed according to the 10X Genomics 5’ single cell user guide.

Three NextSeq runs and one MiSeq runs were performed and the standard 10X Genomics cellranger (version 3.0.2.) pipeline was used to extract FASTQ files and perform data processing. Sequenced reads were aligned to the human GRCh38 reference sequence provided by 10X Genomics (refdata-cellranger-GRCh38 3.0.0.0).

Seurat (v3.1.1) was used for the downstream analysis following the vignette for the analysis of an integrated dataset. Briefly, both samples were first filtered for cells containing at least 200 and less than 4000 features as well as a mitochondrial and ribosomal gene content less than 15% and 25%, respectively. Samples were normalized and variable genes identified using the FindVariableFeatures function with nfeatures limited to 2000. Both samples were integrated, scaled and principal components were calculated. Clustering was performed using Uniform Manifold Approximation and Projection (UMAP) with the first 20 dimensions and a resolution of 0.5. Conserved markers between each cluster for both samples were identified using the FindConservedMarker function and cluster identities were assigned based on previously published marker genes in combination with the web-based tool Enrichr (https://amp.pharm.mssm.edu/Enrichr/). Differentially expressed genes of the no GvHD vs. pre-GvHD sample were calculated using the FindMarker function. The R package EnhancedVolcano was used to generate the volcano plots based on the differential gene expression within a cluster.

**Table 2:** scRNAseq sample output metrics summary

| Sample | Estimated Number of Cells | Mean Reads per Cell | Median Genes per Cell | Sequencing Saturation | Total Genes Detected | Median UMI Counts per Cell |
| --- | --- | --- | --- | --- | --- | --- |
| No GvHD | 6,830 | 66,564 | 1,593 | 84.10% | 19,734 | 4,289 |
| Pre-GvHD | 5,219 | 62,829 | 1,468 | 83.20% | 18,625 | 4,272 |

### Statistics

Statistical analyses were performed using Prism (GraphPad Software, USA). Error bars on bar graphs represent mean ± SEM. Experimental groups were compared using a mixed-effects model with Sidak’s multiple comparison test or two-way analysis of variance followed by Tukey’s multiple comparison test. A p-value <0.05 was considered statistically significant, *p < 0.05, **p < 0.01, and ***p < 0.001.

## Results

### Intracellular metabolic screen identifies high intracellular lactate within alloreactive CD4^+^ T cells

We previously demonstrated infiltration of liver, gut, skin, and spleen by donor-derived alloreactive CD4^+^ T cells in the B10.D2 into BALB/c mouse model^23^. These cells were predominantly of the effector memory (T_em_) phenotype (CD44^high^CCR7^-^) at day 14, when mice still appear largely asymptomatic (Figure 1A), i.e. are gaining weight (Figure 1B) and show no overt signs of disease (Figure 1C). Meanwhile, CD4^+^ T_em_ cells continue to predominate in the target organs when animals manifest measurable clinical signs of disease^23^. Thus, we sought to elucidate the metabolic features of CD4^+^ T cell subsets on day 14, a presymptomatic time point, to gain insight into GvHD pathogenesis, and potentially exploit metabolic properties of infiltrating T cells for non-invasive early detection of GvHD.

**Figure 1.**
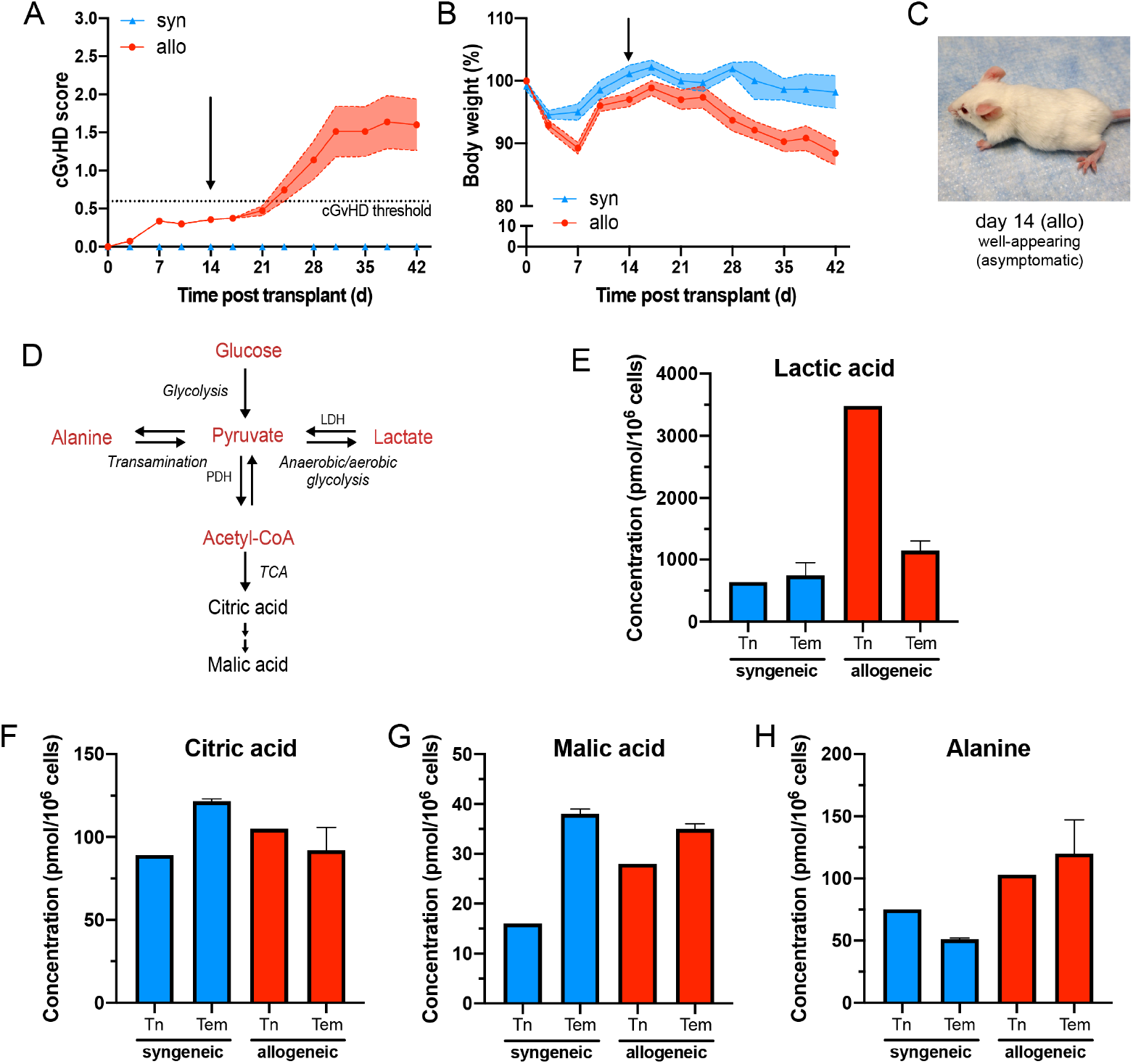
Mass spectrometry screen of intracellular metabolites in CD4^+^ T cell subsets early in cGvHD reveals increased aerobic glycolysis in allogeneic T_em_ cells. Representative example of the clinical score **(A)** and body weight **(B)** changes over time in the B10.D2 into BALB/c GvHD model, n= 5 (syn), n = 16 (allo). **C)** At day 14, mice appear asymptomatic with the exception of erythematous ears. **D)** Schematic indicating the potential fate of pyruvate by either being converted to lactate (anaerobic/aerobic glycolysis), acetyl-CoA (TCA cycle) or alanine (transamination). **E-H)** Intracellular concentrations of metabolites: lactate (E), citric acid (F), malic acid (G) and alanine (H). Single cell suspensions were generated from pooled, freshly harvested spleens of syn (n = 19) and allo (n = 22) HSCT recipients on day 14. The cells underwent positive selection using CD4 microbeads. FACS-purified T cell subsets were collected, and enzyme activity quenched with methanol. All samples were supplemented with an internal standard solution and relative quantification was carried out using CE-TOF mass spectrometry. Cells were pooled from two independent HSCTs. Each sample for MS analysis contained ~3 million cells; n= 1 for syn/allo T_n_ samples, n = 2 for syngeneic T_em_ and n = 3 for allogeneic T_em_. Data is represented as mean + SEM, as appropriate.

Activated CD4^+^ and CD8^+^T cells have been shown to have a distinct metabolic profile, including elevated glycolysis in settings of cancer and autoimmunity^44–46^. We therefore focused on metabolites generated from pyruvate, the substrate precursor situated at the crossroads between aerobic glycolysis, the tricarboxylic acid (TCA) cycle and amino acid metabolism (Figure 1D). We performed an intracellular metabolomic screen using FACS-purified CD4^+^T cell subsets isolated from pooled spleen samples of allogeneic graft recipients on day 14 and compared them to phenotypic counterparts isolated from syngeneic recipients that were subjected to the same HSCT conditioning regimen.

We found that phenotypically naïve CD4^+^ T cells (T_n_, CD44^low^CCR7^+^) in allogeneic HSCT recipients had 3-4-fold higher intracellular levels of lactate compared to allogeneic T_em_ as well as CD4^+^ T cells extracted from syngeneic recipients (Figure 1E). This indicates that while these cells still express CCR7 and have low CD44 expression they are likely antigen stimulated. Allogeneic CD4^+^ T_em_ showed a 50% higher intracellular lactate compared to their syngeneic counterparts. In contrast, citric acid and malic acid, intermediates of the TCA cycle, were more abundant in syngeneic T_em_ cells (Figure 1F,G). Allogeneic CD4^+^ T cells showed no differences between T_n_ and T_em_ with regard to citric acid, while malic acid levels were increased in both subsets compared to syngeneic T_n_ cells. No clear difference in intracellular alanine between syngeneic T_n_ or T_em_ was detected, while alanine levels in both allogeneic T cell subsets were slightly higher compared with syngeneic subsets (Figure 1H). Based on these results, increased conversion of pyruvate to lactate appeared to be a distinguishing metabolic feature of allogeneic CD4^+^ T cells.

Next, we sought to evaluate *ex vivo* metabolism of CD4^+^T cell subsets using a metabolic flux assay that measures the extracellular acidification rate (ECAR) and the oxygen consumption rate (OCR).

### CD4^+^ T_em_ cells extracted from allogeneic HSCT recipients display a high ECAR

We FACS purified CD4^+^ T_n_ and T_em_ cells from spleens of allogeneic and syngeneic HSCT recipients on day 14 and measured their ECAR and OCR, surrogate measures for glycolysis and oxidative phosphorylation, respectively (Figure 2A,B). Allogeneic CD4^+^Tem cells were distinguished from other subsets by the highest basal ECAR (Figure 2B). While both syngeneic and allogeneic T_em_ cells showed elevated basal glycolysis compared to T_n_, this difference was more pronounced between allogeneic T_em_ and T_n_ cells with the highest basal glycolysis observed in allogeneic T_em_ (Figure 2C). Similarly, the glycolytic capacity of allogeneic T_em_ cells was higher compared to the other subsets (Figure 2D). Syngeneic T_em_ had a lower basal respiration and basal glycolysis than allogeneic T_em_. Based on the high extracellular lactate export (i.e. ECAR) and the low intracellular lactate levels, we hypothesized that upon differentiation from a naïve surface phenotype, T_em_ cells continue to rely on glycolysis and produce lactate, but do not accumulate lactate intracellularly due to an increased extracellular transport.

**Figure 2.**
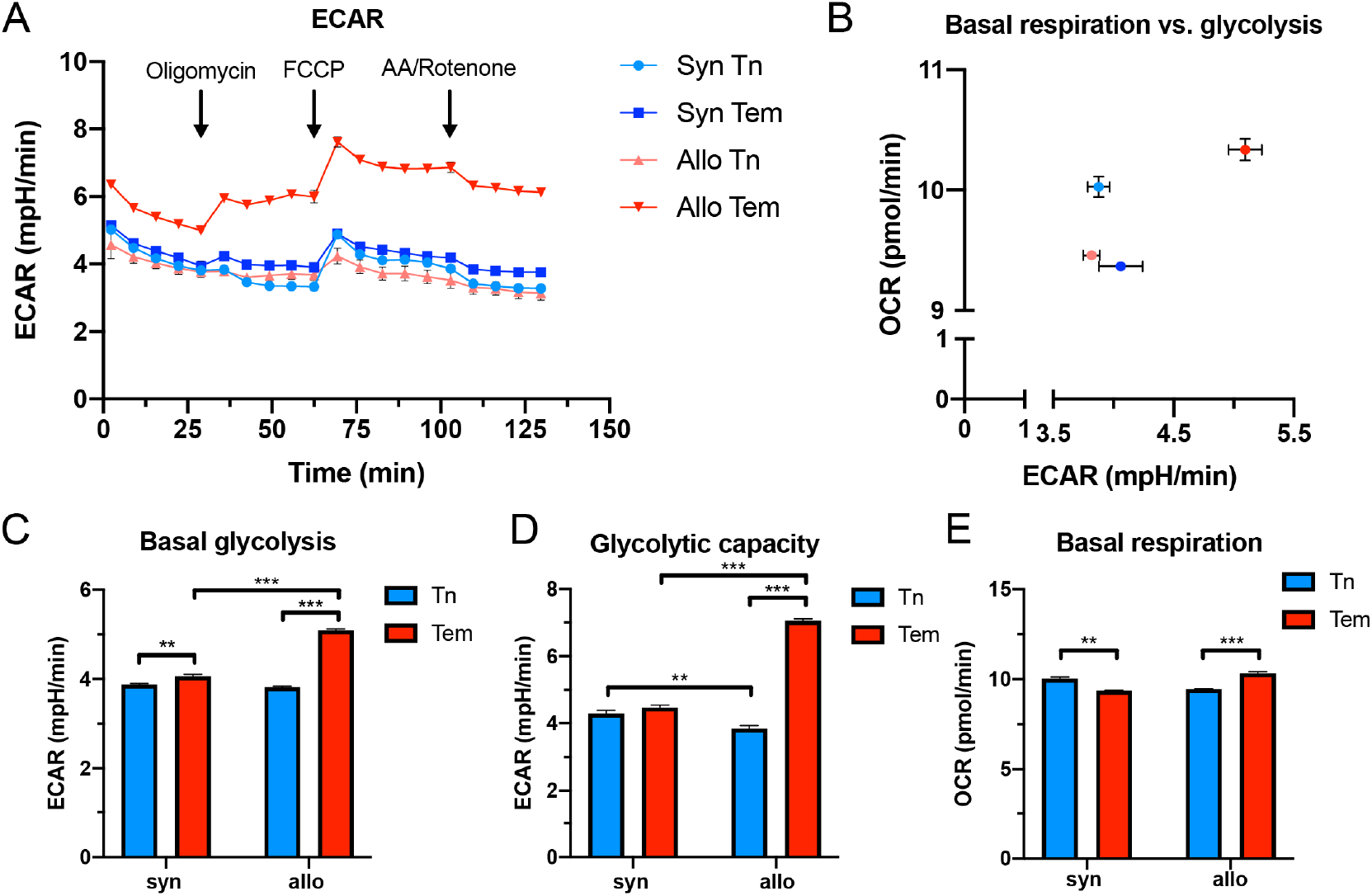
CD4^+^ T_em_ cells exhibit a higher extracellular acidification rate compared to CD4^+^ T_n_ or syngeneic CD4^+^ T_em_ in cGvHD. **A)** Single cell suspensions were generated from pooled harvested spleens of syngeneic (n = 5 mice) and allogeneic (n = 9 mice) HSCT recipients on day 14. The cells underwent positive selection using CD4 magnetic microbeads, followed by FACS. FACS-purified T cells were plated using CellTak at a seeding density of 100,000 live sorted cells per well and treated with oligomycin (1 μM), FCCP (0.5 μM) and antimycin A/Rotenone (1 μM) at indicated timepoints. The number of technical replicates equals n = 8-32 wells/subtype, data are representative of 2 independent experiments. **B)** Basal glycolysis versus basal respiration. **C)** Basal glycolysis based on the mean of the two timepoints before oligomycin injection. **D)** Glycolytic capacity measured after FCCP addition (mean of four timepoints). **E)** Basal respiration rate based on the mean of the three timepoints before oligomycin injection. Data is represented as mean + SEM, statistical testing using a two-way ANOVA, *=p<0.05.

### CD4^+^ T_em_ cells upregulate transcription of glycolytic enzymes and transporters

To gain additional insight into the metabolic properties of CD4^+^ subsets, we performed RNA sequencing on splenic allogeneic T_em_ and T_n_ cells to compare their expression of key metabolic enzymes in glycolysis and the TCA cycle (Figure 3). Consistent with our intracellular and *ex vivo* metabolic data, higher transcriptional levels of multiple glycolytic enzymes were detected in T_em_ versus T_n_ cells. Specifically, transcripts for key regulatory glycolytic enzymes such as hexokinase 2 (*Hk2*), which catalyzes the commitment step of glycolysis, glyceraldehyde 3-phosphate dehydrogenase (*Gapdh*) and pyruvate kinase (*Pkm*), which serves as a rate-limiting step, were expressed at higher levels in allogeneic CD4^+^ T_em_ compared to T_n_ cells (Figure 3). Along with increased transcriptional expression of glycolytic enzymes, glucose transporters *Slc2a1* and *Slc2a3* (Glut1 and Glut3, respectively) were also upregulated in the CD4^+^ T_em_ subset suggesting increased glucose import into these cells. In addition, transcripts for the lactate transporter *Slcl6a1* (Mct1) were more highly expressed in T_em_ compared to T_n_ cells. This may explain the observed decreased levels of intracellular lactate concurrent with the higher ECAR in allogeneic T_em_ compared to T_n_.

**Figure 3.**
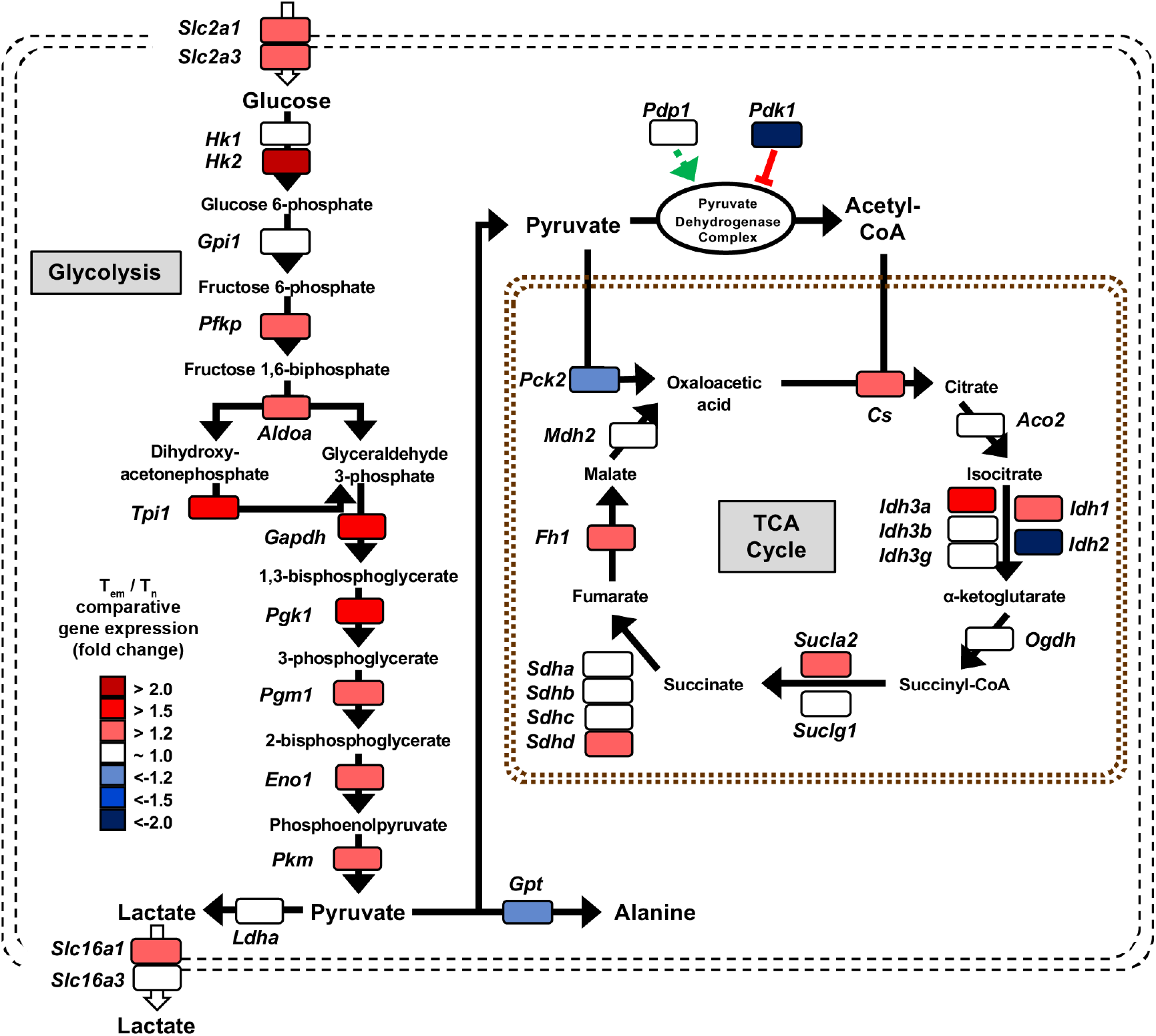
Bulk RNA sequencing of allogeneic CD4^+^ T_n_ and T_em_ cells indicates an overall upregulation of glycolytic enzymes in alloreactive T_em_ cells. FACS-purified allogeneic T_em_ and T_n_ cells from the spleen (> 0.5 million cells per sample) were pooled from multiple study animals on day 14. RNA was extracted from pooled samples of each cell type (n = 1 for T_n_, n = 3 for T_em_) and the library was generated using the HyperPrep RNA-Seq Kit. All samples were sequenced on a HiSeq4000 (Illumina) and the reads were trimmed, mapped to the reference genome and normalized to the library size as counts per million. The shaded squares indicate an increased (red) or decreased (blue) expression in T_em_ cells over T_n_ cells. *Slc2a1*-solute carrier family 2 member 1 (GLUT1); *Slc2a3-* solute carrier family 2 member 3 (GLUT3); *Hk1*-hexokinase 1; *Hk2*-hexokinase 2; Gpi1-glucose phosphate isomerase 1; *Pfkp*-phosphofructokinase; ldoa-fructose-bisphosphate aldolase A; *Gapdh*-glyceraldehyde-3-phosphate dehydrogenase; *Pgkl*-phosphoglycerate kinase 1; *Pgml*-phosphoglucomutase1; *Enol*-enolase 1; *Pkm*-pyruvate kinase; *Ldha-* lactose dehydrogenase A; *Slc16a1*-solute carrier family member 16 (MCT1); *Slc16a3*-solute carrier family 2 member 1 (MCT4); *Gpt*-glutamate pyruvate transaminase; *Pdp1*-pyruvate dehydrogenase phosphatase 1; *Pdk1*-pyruvate dehydrogenase kinase 1; *Pck2*-phosphoenolpyruvate carboxykinase 2; *Cs*-citrate synthase; *Aco2*-aconitase 2; *Idh*-Isocitrate dehydrogenase 1; *Ogdh*-oxoglutarate dehydrogenase; *Sucla*-succinyl-CoA ligase; *Sudh*-succinate dehydrogenase; *Fh*-fumarate hydratase; *Mdh*-malate dehydrogenase; *Acyl*-acyl-CoA synthetase; *Acaca*-acetyl-CoA carboxylase.

Taken together and in line with intracellular and *ex vivo* metabolic data, RNA sequencing of CD4^+^ T cell subsets indicates that glycolysis is upregulated upon T cell transition from naïve to effector memory phenotype.

### Hyperpolarized [1-^13^C]pyruvate MRI demonstrates high conversion of pyruvate to lactate within the liver

Since highly glycolytic activated T cells infiltrate target organs, we hypothesized that these metabolic changes could be exploited to visualize affected organs differentiating them from unaffected tissues or syngeneic recipients. To test this, we performed ^13^C-hyperpolarized MRI over the post-HSCT course. As stated, the method tracks ^13^C-labeled hyperpolarized pyruvate and its ^13^C-labeled metabolites in tissue (Figure 4A). We performed serial ^13^C-MRI scans on mice that underwent syngeneic or allogeneic HSCT starting on post-HSCT day 7 and continued through day 28 at regularly scheduled intervals. The liver was first defined anatomically as a region of interest and, immediately following the infusion of the hyperpolarized [1-^13^C] pyruvic acid (12 μL/g), the peaks for [1-^13^C] pyruvate and its metabolites [1-^13^C] alanine and [1-^13^C] lactate were measured at approximately 171, 177, and 183 ppm, respectively (Figure 4B). On day 14, we observed a significantly increased liver lactate/pyruvate ratio in allogeneic HSCT recipients compared to syngeneic controls (mean 1.65 vs. 0.49, p=0.01) indicating higher rates of LDH activity and increased generation of lactate in the livers of allogenic HSCT recipients (Figure 4C,D). This difference was only observed on day 14, suggesting that glycolytic activity in the GvHD target organ was dynamic and transient (Figure 4D). Importantly, increased conversion of pyruvate to lactate was detectable in the liver before animals developed clinical signs of GvHD. We inferred that the elevated lactate conversion in the liver was at least in part due to high rates of lactate production and extracellular secretion by infiltrating alloreactive CD4^+^T_em_ cells. To test whether elevated lactate production in the liver leads to concurrent systemic lactate elevation, we collected serial plasma samples over the post-HSCT course to quantify the plasma lactate concentration (Figure 4E). Interestingly, we observed that the plasma lactate levels peaked on day 7 and were significantly higher in allogeneic than in syngeneic animals, whereas other timepoints showed no significant differences between cohorts (Figure 4E). Concurrently we assessed the number of circulating alloreactive CD4^+^ T cells and found significantly higher numbers of circulating T_em_ cells in allogeneic compared to syngeneic recipients on day 7 and 14, with a higher number on day 7 (Supplemental Figure 1).

**Figure 4.**
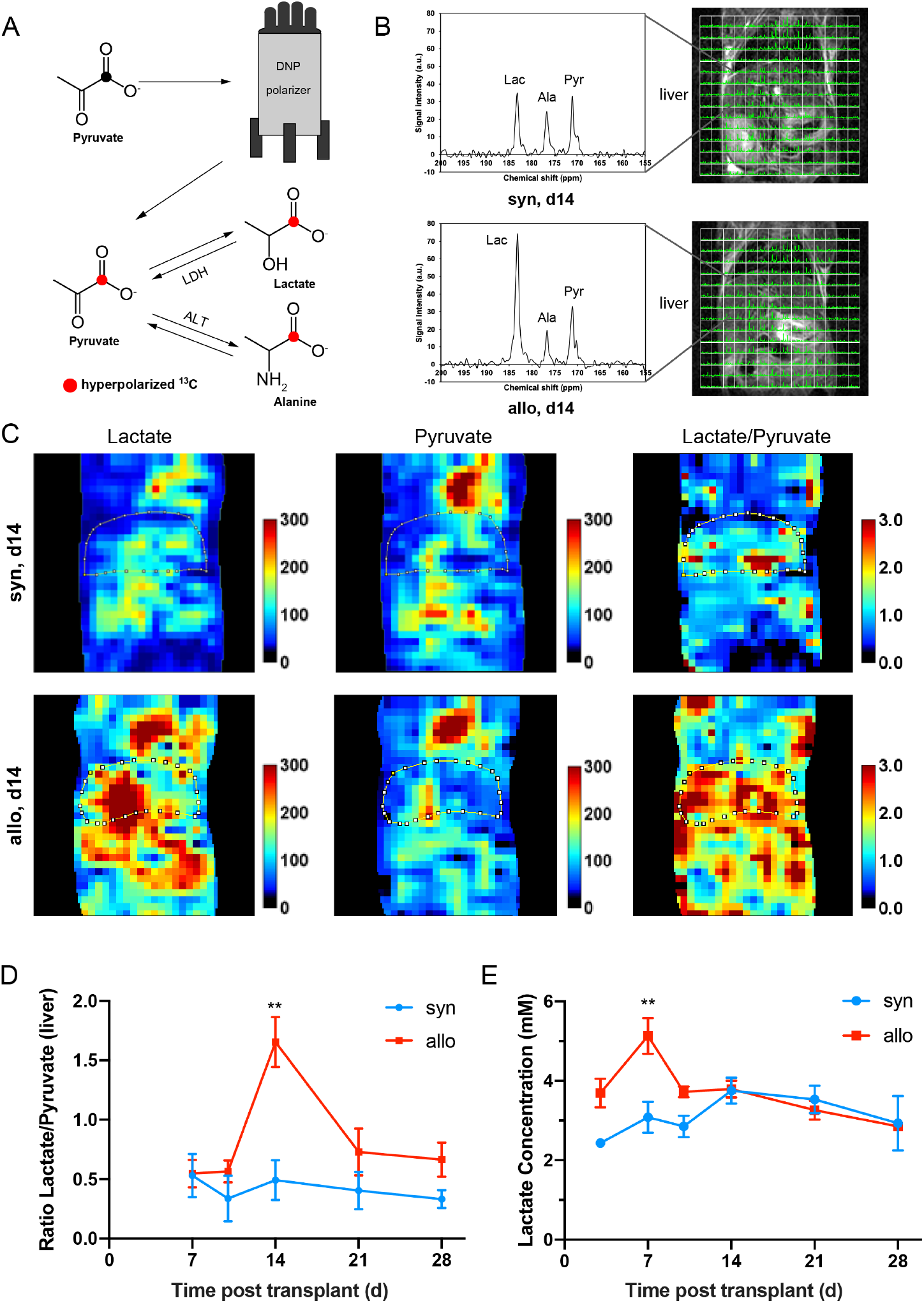
Hyperpolarized ^13^C-pyruvate *in vivo* MRI performed over the post-syngeneic and allogeneic HSCT course. [1-^13^C]pyruvic acid (30 μL), containing 15 mM OX063 and 2.5 mM gadolinium chelate, was hyperpolarized using the Hypersense DNP Polarizer (Oxford Instruments, Abingdon, UK). After hyperpolarization was achieved, the sample was dissolved in 4.5 mL of heated alkaline buffer, i.e. 40 mM 4-(2-hydroxyethyl)-1 piperazineethanesulfonic acid, 30 mM of NaCl, and 100 mg/L of EDTA. The hyperpolarized [1-^13^C] pyruvate solution (96 mM) was then administered through an intravenous tail vein catheter. ^13^C MRI studies were performed on a 3T MR Solutions MRI using a custom ^13^C-^1^H saddle coil for spectroscopic and anatomical imaging. **A)** Schematic representation of the pyruvate to lactate conversion with the hyperpolarized ^13^C indicated. **B)** Representative coronal anatomical MRI images overlaid with the ^13^C MRSI spectra for each voxel and an exemplary ^13^C spectrum indicating the lactate, alanine and pyruvate peaks shown for a syngeneic (top) and allogeneic (bottom) animal at day 14. **C)** MRSI images of syn (top) and allo (bottom) at day 14 indicating the signal intensity for lactate, pyruvate as well as the combined lactate/pyruvate ratio. A region of interest was drawn around the liver based on the anatomical MRI which was then used to quantify the signal intensity for each peak. **D)** Lactate/Pyruvate signal intensity in the liver over time after HSCT in syngeneic and allogeneic animals, n= 3-6 animals per group and timepoint, ** = p<0.01 in a mixed-effect analysis with Sidak’s multiple comparison test. **E)** Mouse plasma was collected via mandibular bleed at indicated timepoints in sodium fluoride/EDTA coated microtubes. After centrifugation, an internal standard was added, and the lactate levels were quantified using UPLC-mass spectrometry, n = 4-17 per group and timepoint, ** = p<0.01 in a mixed-effect analysis with Sidak’s multiple comparison test. All data is shown as mean ± SEM.

### Single-cell RNA sequencing (scRNAseq) of human peripheral blood mononuclear cells (PBMCs) indicates increased transcription of glycolytic enzymes before the onset of aGvHD

To evaluate whether metabolic changes within CD4^+^T cells observed in the pre-clinical model are also present in patients, we performed scRNAseq on human PBMCs obtained from patients that underwent T cell-replete AHSCT. We retrospectively screened annotated clinical data from a GvHD clinical study performed at the National Cancer Institute, USA (NCT00520130). Our goal was to perform scRNAseq of PBMCs collected on day 30 (standard collection time point for the trial) from patients that met the following criteria: 1) no evidence of acute or chronic GvHD (up to 1 year post-HSCT) or 2) those that developed acute GvHD shortly after the day 30 collection date, but were asymptomatic at the time of PBMC collection. One patient for each criterion was successfully identified (for details, see Supplemental Methods). Both patients underwent the same reduced intensity conditioning regimen and received an 8/8-matched graft from an unrelated donor. The “pre-GvHD” patient became symptomatic five days after and was diagnosed with aGvHD seven days post PBMC collection.

We analyzed the gene expression profile of the “no-GvHD” and “pre-GvHD” PBMCs described above using the R package Seurat^47,48^. A total of 5969 and 4297 cells were analyzed for the “no GvHD” and “pre-GvHD” patient, respectively (Figure 5A). UMAP clustering generated 14 distinct clusters after integration of both samples and their identity was assigned using known cellular markers published in previous scRNAseq datasets (Figure 5B,C). Both samples consisted mainly of monocytes (~60-70%) as well as dendritic cells (DC), natural killer (NK) and T cells. Small numbers of contaminating platelets and erythrocytes were also observed (Figure 5D,E). T cells further clustered into separate CD4^+^ and CD8^+^ as well as a mixed CD4/CD8 cluster that was strongly associated with gene expression indicating proliferation, e.g. MKI67 (designated as T_proliferating, Supplemental Figure 2A). Comparing the gene expression of the CD4^+^ and T_proliferating clusters between both patients showed increased expression of genes associated with T cell activation, i.e. *CD69, GADD45B* as well as *JUN* and *FOS* (Figure 5F). In addition, *SLC2A3* expression, the gene for the glucose transporter GLUT-3 was significantly increased in both CD4^+^T cell containing clusters of the “pre-GvHD” patient sample.

**Figure 5.**
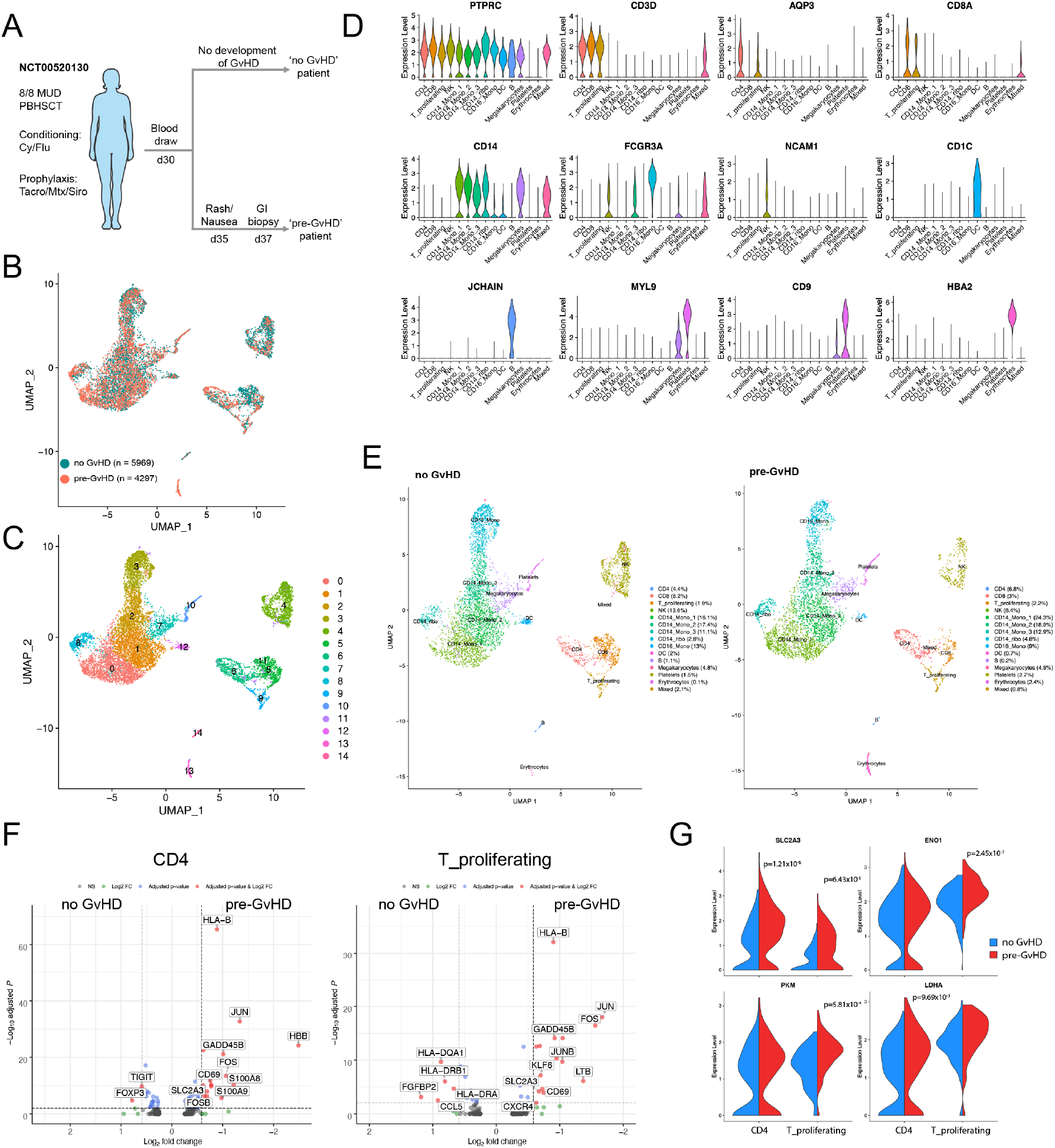
Single-cell RNA sequencing of AHSCT patient-derived PBMCs. **A)** Diagram depicting sample selection. PBMCs of two patients, one of which developed GvHD shortly after sample collection (pre-GvHD) and one who did not (no GvHD) were thawed, red blood cells were lysed, and cell viability was assessed. Single-cell preparation was performed using the Chromium Next GEM Single Cell 5’ Library & Gel Bead Kit (10X Genomics) with a recovery target of 6,000 cells per sample according to the manufacturer’s instructions. The standard 10X Genomics *cellranger* pipeline was used to extract Fastq files and to perform data processing. Sequenced reads were aligned to the human GRCh38 reference sequence provided by 10X Genomics. Clustering and visualization were performed in R using the Seurat package (v3.1.1) with integrated datasets. **B)** UMAP visualization of both samples and the total cell number that went into the analysis. **C)** Clustering result of the integrated data set yielding 14 separate clusters. **D)** Gene expression of known PBMC markers used to assign cluster identities. **E)** UMAP visualization of assigned clusters for both samples with the cluster frequency indicated for each cell type. **F)** Volcano plot of the differential gene expression within one cluster comparing the no GvHD vs. pre-GvHD sample for the CD4 cluster (left) and T_proliferating cluster (right). Genes with a fold-change >1.5 and adjusted p-value <0.01 are highlighted in red. **G)** Violin plot visualizing the gene expression level of key glycolysis enzymes for the CD4 and T_proliferating cluster. Significant adjusted p-values based on the MAST-DE test are indicated in the plot.

While CD8^+^T cells from the “pre-GvHD” sample also showed an increase in activation-related genes, *SLC2A3* transcription did not appear to be increased (Supplemental Figure 2B). The expression level of key genes within the glycolysis pathway for the two CD4^+^T cell containing clusters indicated that several genes, i.e. *SLC2A3, PKM, ENO1* and *LDHA,* tended to have a small, but significantly increased expression in the “pre-GvHD” patient sample (Figure 5G). This was only observed for the CD4^+^ and T_proliferating clusters but not CD8^+^T cells (Supplemental Figure 2C).

## Discussion

In this manuscript we describe a metabolic MRI approach that detects target organ involvement ahead of overt symptoms in a pre-clinical chronic GvHD model. The early post-HSCT window that we visualized *in vivo* is characterized by inflammatory changes that set the course for subsequent fulminant cGvHD manifestations^23^. We also found concurrent increased glycolytic metabolism and transcription of glycolytic enzymes within alloreactive CD4^+^ T cells. Furthermore, single cell RNA sequencing of circulating CD4^+^ T cells from patients undergoing HSCT suggests increased transcription of glycolytic genes prior to onset of overt acute GvHD clinical symptoms.

Consistent with previous work on CD4^+^ T cell metabolism in mouse models of GvHD, we observed a glycolytic phenotype in allogeneic CD4^+^ T_em_ cells. Specifically, an upregulation of glucose and lactate transporters in CD4^+^ T cells in acute GvHD models has been previously described^49,50^, and our research reports the glucose transporters *Glut1* and *Glut3* to be upregulated early in the context of chronic GvHD. However, despite increased lactate excretion by CD4^+^Tem cells, we did not observe a further upregulation of *Ldha* transcription. This likely stems from the fact that we compared allogeneic CD4^+^T_em_ to T_n_ cells while previously bulk CD4^+^T cells from allogeneic and syngeneic cohorts were compared^49^. Since allogeneic T_n_ cells in our study already had high levels of intracellular lactate, indicating high activity of LDHA, the latter is likely sustained through the phenotypic transition to T_em_ without further upregulation or regulated post-transcriptionally, e.g. via phosphorylation^51^.

Regarding the role of oxidative phosphorylation in T cells during GvHD, some studies have demonstrated a parallel increase of both glycolysis and oxidative phosphorylation early posttransplant in acute GvHD models^52^ while others highlight glycolysis as the dominant metabolic pathway^49,50^. Our data also indicate a less dominant role for oxidative phosphorylation compared to glycolysis in GvHD as demonstrated by small differences in OCR between syngeneic and allogeneic T cells and a less uniform upregulation of TCA cycle gene expression within allogeneic CD4^+^ T cell subsets.

Importantly, our study indicates that T cell metabolism is dynamic over the course of GvHD. While we detected high rates of pyruvate to lactate conversion in the liver on day 14, later time points showed no differences, pointing to further metabolic shifts upon disease progression. Prior studies of T cell metabolism in GvHD identified glycolysis as a critical metabolic pathway for T cell activation and antigenic priming during disease initiation but the reliance on glycolysis can change during later stages of T cell activation and differentiation^53^. Several other metabolic pathways pertinent to T cell function in GvHD have already been identified, i.e. fatty acid oxidation, glutaminolysis and others^49,54,55^. Since a wide range of metabolites are amenable to ^13^C-metabolic tracing *in vivo*^56^, further characterization of immune metabolism over the entire course of GvHD is prudent and may elucidate unique features of early, ongoing and well-established GvHD *in vivo*^53^.

Whether increased glycolytic flux observed by metabolic MRI is solely attributable to T cells infiltrating the liver is not yet known. Other immune populations present in the liver, such as macrophages and dendritic cells, have been described to upregulate glycolysis in response to inflammatory stimuli^57,58^ and non-hematopoietic cells, such as hepatocytes, could potentially experience a perturbed glucose metabolism in the setting of an immune attack^59^. However, in support of our hypothesis that T cell infiltration is a major contributor to the observed MRI liver signal, Pektor *et al*. have shown that alloreactive T cells are most abundant in the liver in a humanized mouse model of GvHD at a similar time point when assessed via whole body PET imaging using a ^89^Zr-labeled anti-human CD3 antibody^60^. When another T cell specific PET radiotracer was tested in an acute GvHD model, high background uptake in the liver precluded the evaluation of T cell trafficking despite previous studies confirming trafficking to the liver using bioluminescence imaging^61^ and other techniques^23,62^. Interestingly, our images of the GI tract, another key GvHD target tissue infiltrated by alloreactive T cells on day 14^23^, suggest increased glycolytic activity on day 14, albeit the quantification was not performed due to the gut anatomy being less conducive than liver for accurately mapping regions of interest to correlate with metabolic tracing. Additionally, we did not observe increased glycolytic flux in the liver in the setting of syngeneic HSCT, which is in line with FDG-PET GvHD imaging where FDG uptake correlated with concurrent target organ (colon) infiltration by EGFP^+^ donor lymphocytes and was similarly not observed in the syngeneic HSCT setting or in the absence of T cells in the tissue^63^.

Our approach is safe to translate to the clinic because unlike FDG-PET it lacks exposure to ionizing radiation and thus can be performed serially in the same patient making it ideal for evaluating the dynamic post-transplant course. Wide clinical implementation of hyperpolarized metabolic imaging may be currently limited by the logistics and cost of equipment, but active clinical studies testing methodological modifications are currently being conducted and could facilitate wider application of metabolic MRI^30,64^. Furthermore, recent improvements in data processing have enabled omission of hyperpolarization for metabolic imaging and could be tested in the setting of GvHD^65,66^.

While metabolic imaging is safe, its clinical utility in the post-HSCT setting may be limited by the potential lack of specificity for GvHD. T cell activation is likely to play a role in several processes that would be important to distinguish from GvHD. Infection, graft rejection, engraftment, relapse, and early GvHD may have similar radiographic features. However, using this imaging modality in combination with established routine clinical testing may help distinguish GvHD from the other above mentioned immune driven complications of HSCT. Additionally, serial plasma lactate sampling, which is relatively easy and inexpensive to collect and analyze, could be helpful in identifying an optimal time for performing metabolic imaging. The observed rise in plasma lactate prior to target organ infiltration could be related to the high number of circulating activated T cells producing and exporting lactate^67^. Prospective single-cell RNA sequencing of PBMCs early post HSCT may elucidate changes in the T cell compartment prior to disease onset. We saw transcriptomic evidence of increased T cell activation in the PBMC sample of a patient that developed acute GvHD a week after the analyzed time point. The observed increased expression of *SLC2A3* (GLUT3) was also seen in our pre-clinical model and by others at early GvHD time points, indicating that further evaluation of glycolytic metabolism, including via metabolic imaging, may be useful in patients at risk for GvHD^49^. A larger sample size and a timecourse evaluation of immunometabolic transcriptome in the post-HSCT setting is warranted given our findings.

Finally, therapeutic targeting of upregulated glycolysis is a promising strategy for modulating overactive immunity that characterizes GvHD and other T cell-mediated conditions. Multiple steps in the glycolytic pathway have already been inhibited to augment T cell function in auto- and alloimmune diseases^49,50,55,68,69^ and ^13^C-pyruvate imaging can be used to detect responses to glycolytic inhibitors^70,71^. Non-invasive and non-radioactive approaches to guide clinical decisions are needed, and this methodology offers a promising new tool for the post-HSCT setting. It has the potential to not only discern the presence or absence of GvHD in a non-invasive manner, but to also illuminate *in vivo* metabolic shifts that are amenable to therapeutic targeting.

## Acknowledgements

We are grateful to Devorah Gallardo for expert assistance during animal experiments. We acknowledge Kevin Hu for helping with the collection of samples for the intracellular metabolomic profile. We are grateful to Veena Kapoor and William Telford of the ETIB Flow Cytometry Core facility for performing FACS purifications. We are grateful to Jeremy Rose of ETIB Clinical Core for help with selecting clinical samples from NCT00520130. We thank the National Institutes of Health (NIH) Intramural Sequencing Center (NISC) and National Cancer Institute (NCI) Single Cell Analysis Facility (SCAF) for their technical support. This was work was supported by intramural funding from NCI, NIH. Support from CCR Single Cell Analysis Facility was funded by FNLCR Contract HHSN261200800001E. Sequencing was performed with the CCR Genomics Core. This work utilized the computational resources of the NIH HPC Biowulf cluster (http://hpc.nih.gov).

## Author contributions

NPB conceived, designed and supervised the study. NM, BO, DEF and NPB performed the intracellular metabolic screen. CS, NM and NPB carried out the metabolic flux assays. CJR, TJM, KW, NM, JCA and NPB generated and analyzed the RNA sequencing data. KS carried out the ^13^C-hyperpolarized MRI. KW, JCA, and NPB collected and DEF quantified the plasma lactate levels. KW and JCA quantified the circulating T cells. KW, JCA and NPB generated samples for scRNAseq. JCA analyzed the scRNAseq data. SP provided the patient samples. MCK, WML, REG provided critical instruments and reagents. JCA and NPB wrote the manuscript. All authors contributed to the editing and approved the final manuscript.

## Supplemental Figures

**Supplemental Figure 1.**
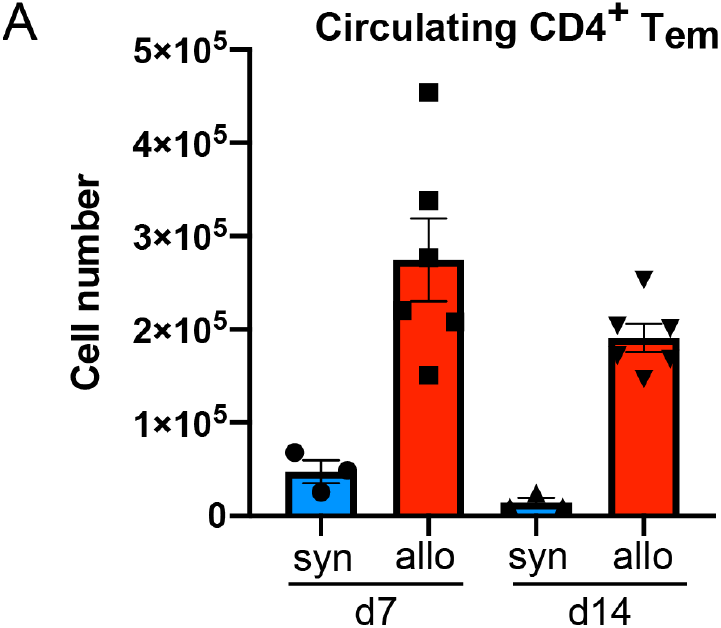
Circulating T_em_ cells at early timepoints post-HSCT. To quantify circulating CD4^+^ T_n_ and T_em_, blood was drawn via cardiac puncture into heparin-coated tubes, red blood cells were lysed, and the remaining cells were stained with antibodies against CD4, CD44, CCR7, CD25, FOXP3 and CD69 (T_em_) defined as CD4^+^, FOXP3^-^,CCR7^-^, CD44^high^), n = 3-6, all data shown as mean ± SEM.

**Supplemental Figure 2.**
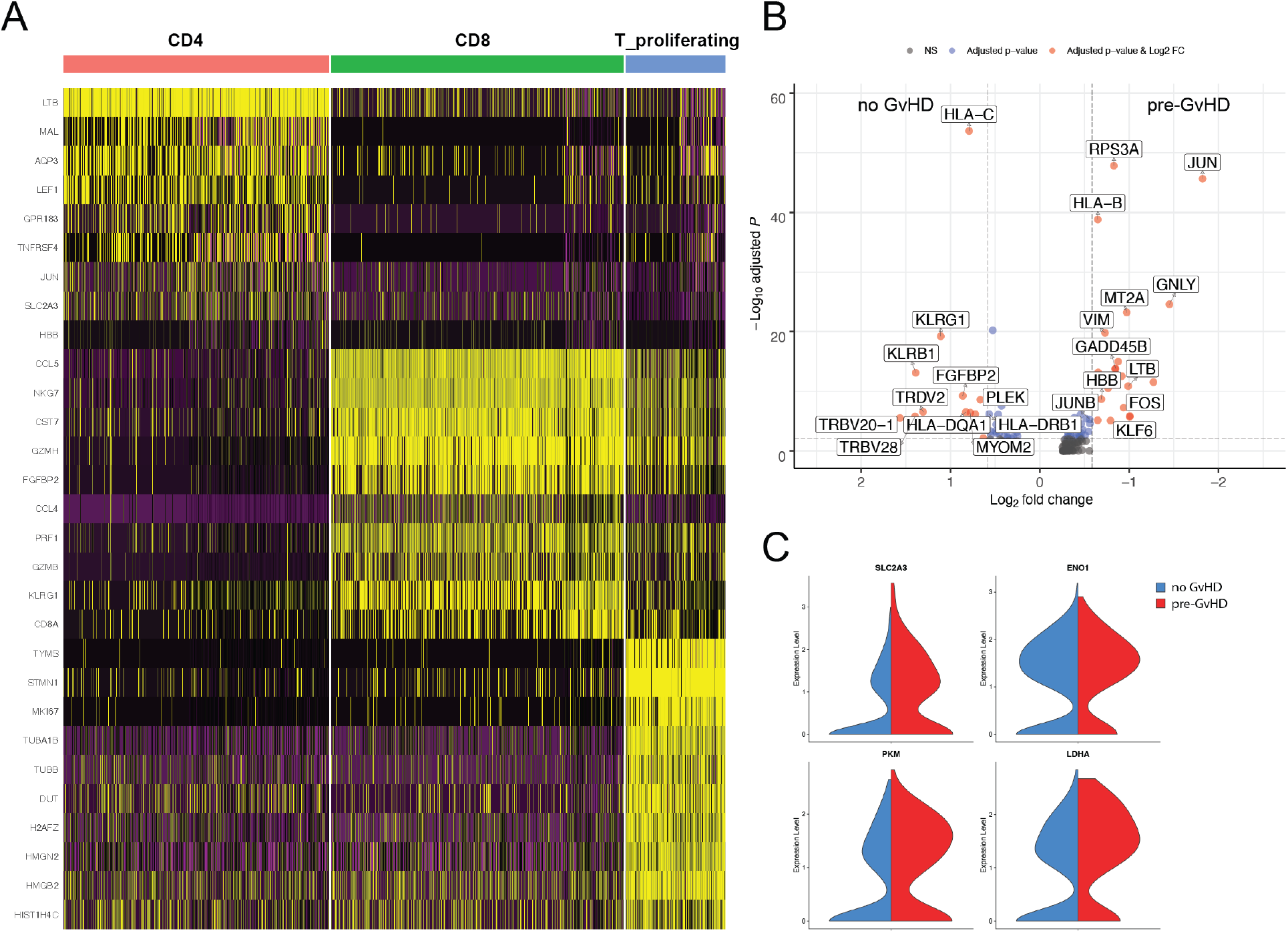
Top 10 conserved markers for T cell-containing clusters and differentially expressed genes in CD8^+^ T cells. **A)** Heatmap illustrating the top 10 conserved genes for each of the three identified T cell clusters. **B)** Volcano plot of the differential gene expression within the CD8 T cell cluster comparing the no GvHD vs. pre-GvHD sample. Genes with a fold-change >1.5 and adjusted p-value <0.01 are highlighted in red. **C)** Violin plots visualizing the gene expression level of key glycolysis enzymes for the CD8 T cell cluster.

